# Acetylcholine esterase of *Drosophila melanogaster*: a laboratory model to explore applications of insecticide susceptibility gene drives

**DOI:** 10.1101/2023.11.10.566664

**Authors:** Natalia Hernandes, Xiaomeng Mollyann Qi, Soumitra Bhide, Courtney Brown, Benjamin J. Camm, Simon W. Baxter, Charles Robin

## Abstract

**BACKGROUND:** One of the proposed applications of gene drives has been to revert pesticide resistant mutations back to the ancestral susceptible state. Insecticides that have become ineffective because of the rise of resistance could have reinvigorated utility and be used to suppress pest populations again, perhaps at lower application doses.

**RESULTS:** We have created a laboratory model for susceptibility gene drives that replaces field-selected resistant variants of the acetylcholine esterase (*Ace*) locus of *Drosophila melanogaster* with ancestral susceptible variants. We constructed a CRISPR/Cas9 homing drive and found that homing occurred in many genetic backgrounds with varying efficiencies. While the drive itself could not be homozygosed, it converted resistant alleles into susceptible ones and produced recessive lethal alleles that could suppress populations. Our studies provided evidence for two distinct classes of Gene Drive Resistance (GDR): rather than being mediated by the conventional Non-Homologous End-joining (NHEJ) pathway, one seemed to involve short homologous repair and the other was defined by genetic background.

Additionally, we used simulations to explore a distinct application of susceptibility drives; the use of chemicals to prevent the spread of synthetic gene drives into protected areas.

**CONCLUSIONS:** Insecticide susceptibility gene drives could be useful tools to control pest insects however problems associated with particularities of the target loci and GDR will need to be overcome for them to be effective. Furthermore, realistic patterns of pest dispersal and high insecticide exposure rates would be required if susceptibility were to be useful as a ‘safety-switch’ to prevent the unwanted spread of gene drives.

## Introduction

Chemical insecticides have been essential tools in modern agricultural practices and have played important roles supressing insect vectors of disease. However, inevitably insecticide resistance evolves so that higher doses or different chemistries are required to prevent and control pest outbreaks (1). Here we explore the suggestion that gene drives could be used to put resistance evolution into reverse and restore genetic susceptibility to an insect population (2–4). Gene drives, which can be synthesized using CRISPR/Cas9 technology, bias inheritance so that genetic elements encoding traits such insecticide susceptibility can be spread through free-living populations(5–7).

There are three reasons why ‘susceptibility gene drives’ might be a good idea. Firstly, a susceptibility gene drive might give an insecticide with desirable properties a new lease of life. For example, phosphine has been widely used to protect stored grain for more than 70 years (8) and has a distinct advantage of leaving no residues on grains and yet it’s utility, and the stored grain industries that depend upon it, are threatened by the rise in phosphine resistance alleles among pest beetles (9). Secondly, the higher the dose that an insecticide is used, the more potent it is as an environmental pollutant. A gene drive that lowers the dose of insecticide required to prevent pest outbreaks could be a boon for the ecosystem. In fact, one could extrapolate this idea to imagine that a gene drive could confer a peculiar sensitivity of a pest insect to an otherwise benign chemical, thereby creating a species-specific control mechanism. Thirdly, an idea that we explore here, is that chemical sensitivity could be used to control the unwanted spread of gene drives (10). Such a sensitivity could be engineered into gene drives generally as a ‘safety-switch’ where the chemical is deployed at population borders or after unintended release.

Methods to control the unwanted spread of gene drives have been at the forefront of recent gene drive research. One approach has been to use gene drives that have a high threshold; they only spread if many individuals in the population have them. This makes them less invasive because if they spread to a new population they will be at a low frequency and will not establish (11). Another approach is to tailor gene drives to local populations by targeting genetic variation that is either private to that population or much more abundant in the targeted population (12). A third approach is the Daisy-Chain design (13), where a succession of elements drive each other but the first element in the chain is not a drive and is entirely dependent on its release frequency. As far as we know, no example of this genetically complex drive has been engineered yet. Recently a fourth idea, that of engineering self-eliminating gene drives has been proposed, in which low frequency recombination events either separate gene drive components (14) or cause gene drive elements to excise over time (15). However, all these proposed methods, rely on the stochastic population processes inherent to the demography of the gene drive population. By engineering chemical sensitivity into gene drives (10) we might have a ‘safety-switch’ that could be used to prevent the drive spreading in time and space. Thus, a chemical sensitivity could be an orthogonal tool that could be deployed by gene drive managers.

Recently, a gene drive conferring susceptibility to pyrethroid and organochlorine insecticides was engineered into *Drosophila melanogaster* (16). First however, a resistant allele of the *para* locus (that encodes a voltage gated sodium channel that is the molecular target of these insecticides), had to be edited into a lab population because such alleles do not occur naturally in this species. The so called ‘knock down resistance (kdr)’ variant (L1014F) is however commonly found in pest species and a serendipitous location of a key CRISPR/Cas9 feature – the PAM site-enabled a strategy where only the resistant allele was cleaved by the CRISPR machinery. This enabled a drive to be engineered that worked on *para* from a distant locus which was driven with high efficiency (∼85% of progeny from drive bearing parents bore the drive allele ; 15). One of the many interesting aspects of this study was that it identified fitness costs of the *kdr* allele in *D. melanogaster* that may explain why DDT resistance in this species is associated with other genes (17,18). The authors also raised the possibility that two distinct susceptibility gene drives could be alternated so that susceptible alleles to insecticide A could spread through a pest population while insecticide B was being used to suppress the population (16).

Here we attempt to generate a second susceptibility gene drive in *Drosophila melanogaster* where naturally occurring organophosphate resistance alleles of the acetylcholine esterase gene (*Ace*) would be reverted to their ancestral susceptible state. Despite *Drosophila melanogaster* not being a pest species, it shows unequivocal signs of adapting to organophosphate insecticides through target site modification (18–20). *Ace* has long been known to be associated with insecticide resistance through focussed analyses (21,22) yet it is also a standout adaptive locus in genome wide scans of selective sweeps (20) and for genome wide association studies seeking the genetic basis of organophosphate resistance (23–25). These studies all point to multiple substitutions of amino acids that line the active site gorge of Ace. The effect of four common amino acid substitutions (I199V, G303A, F368Y and G406A) have been kinetically characterised in baculovirus expressed acetylcholine esterase where their inhibition profile against 17 insecticides was characterized (26). Remarkably, different combinations of the amino acid states maximize inhibition constants to different chemicals.

The natural function of acetylcholine esterase (EC 3.1.1.7) is to rapidly hydrolyse the neurotransmitter acetylcholine so that a synapse can be reset after an activation signal has been relayed (27). Unlike most insects, Cyclorrhaphan flies such as *Drosophila melanogaster* have one rather than two acetylcholine esterase genes (28). Alternate splicing leads to two different protein forms, the PA form which contains a cysteine that forms a disulfide bridge to create dimers that are tethered to the membrane of neurons with a GPI anchor, and the PC form that encodes a monomer of unknown, possibly non-catalytic, function.

Organophosphate insecticides such as malathion, are activated in insects to ‘oxon’ forms (such as malaoxon) that then bind to and inhibit acetylcholine esterase and that quickly leads to the death of the exposed insects (29). Carbamate insecticides, which have a distinct chemistry to organophosphates also target Acetylcholine esterase. Two broad types of ‘target site’ organophosphate resistance are observed; one where point mutations lead to amino acid changes in the active site gorge of acetylcholine esterase that prevent the insecticide covalently binding to the catalytic serine and yet preserve acetylcholine activity (21,29) and the other, which is associated with the increased production of the acetylcholine esterase enzyme through gene copy number variation (30–33).

Here we aim to generate a gene drive that reverts three common amino acid substitutions (27%-38% Drosophila Genetic Reference Panel ; 34) that confer organophosphate resistance, back to their ancestral susceptible forms. Our intention is not to generate a gene drive that propagates the utility of organophosphates and carbamates which are environmental toxins yielding effects on non-target species. Instead, we wished to develop a laboratory model to assess how susceptibility gene drives perform. We set out to explore the merits of various designs and factors affecting their effectivity such as gene drive conversion efficiency.

## Materials and Methods

### 2.1 Plasmids and cloning

DNA oligonucleotides encoding four short guide RNA sequences were chosen to cut in exons 4 and 5 of the *Ace* gene. They were synthesized by a commercial provider and cloned into pCFD5_w plasmid (Addgene #112645) following the protocol of Port and Bullock (35). This plasmid allows the multiplexing of sgRNAs driven off a single U6 RNA polymerase III promoter. The sgRNAs are encoded so that they are interspersed with tRNA sequences that get processed so that independent sgRNAs are liberated. pCFD5 also has an *attB* sequence so that microinjected plasmids integrate at an *attP* landing site of known location in the genome (we used attP40 on the second chromosome). Flies bearing the insertion were crossed to a Cas9 line to generate CRISPR mutations (Supplementary Information Figure S1).

Two mini-*Ace* plasmids both of which had the PA splice form modified so that they were immune to sgRNA-cutting were made by ordering custom gene syntheses from a commercial provider and cloning them into the p*UAS*T plasmid (36). The two plasmids differed by whether they encoded organophosphate resistance variants (PA-R: 199V, 303A, 368Y, 406A) or the ancestral susceptible states (PA-S: I199, G303, F368, G406; Supplementary information S5). The amino acid numbering convention we adopt here is that the start methionine is 1, rather than the first amino acid after the 38 amino acid signal peptide has been removed as is used by Menozzi *et al.* (26). Both were modified with ‘immunity changes’ so that the sgRNAs we were using would not cut them – mainly by changing sequences at synonymous sites. However, we erroneously introduced N338K and H341Q mutations. These changes occurred in helix12 in non-conserved periphery of the Ace protein and the results suggest they did not impact function (see below).

To generate the *Ace* Susceptibility Gene Drive (SGD) we inserted three synthetic fragments into an *Ace* pcfd5 plasmid generated above. This was a multi-step process that had a synthetic fragment for one homology arm (syn2: including a modified exon 4 and some of the targeted intron) and another synthetic element (syn3 that encoded a 3xP3 promoter to express a DsRed marker gene with an SV40 PolyA tail) being placed on one side of the sgRNA cassette. Then a third synthetic element (syn5) containing the other homology arm (including a modified exon 5 and some of the targeted intron) was placed on the other side of the sgRNA cassette (Supplementary Information S1 for sequence details). Both the dsRED and the sgRNAs were designed to be transcribed from the opposite strand to *Ace* to prevent undesired splice conformations and with the intent to minimize transcriptional interference.

The site of insertion within the intron was chosen to be the most divergent region in comparison of homologous regions from other Drosophila species. The homology arms not only encoded the insecticide susceptible form of *Ace* but also had synonymous sites altered to make them immune to sgRNA cutting (as described above for the mini-*Ace* constructs).

We returned to this cloning step twice after the initial attempt. The second time we altered two ‘immunity changes’ to correct the N338K and H341Q mutations. On the third time we removed the dsRED gene (syn3) element altogether which meant that integration of this plasmid into flies had to be determined by PCR.

### 2.2 Drosophila lines

The following Drosophila stocks were obtained from the Bloomington Drosophila Stock Centre: an *Ace* deficiency line (BL7974: w^1118^; Df(3R)Exel8158/TM6B,Tb^1^), an elav-Gal4 line (BL458: P{w{+mW.hs]=GawB}elav^c155^), an nSyb-Gal4 line (BL51635: y^1^ w*; P{+m*]=nSyb-Gal4.S}3, a nos-Cas9 on the X chromosome (BL54591: y^1^ M{w[+mC]=nanos-Cas9.p}ZH-2A w*) and the following DGRP lines (BL28151: Ral-181,BL25177: Ral-304, BL55018: Ral-319, BL25192: Ral-399, BL25198: Ral-555, BL28218: Ral-703, BL28235: Ral-802, BL28238: Ral-808, BL28240: Ral-812, BL28241: Ral-818,BL28249: Ral-850, BL28260: Ral-897). A line with nos-Cas9 on the 2nd chromosome was obtained from the National Institute of Genetics in Japan: NIG-FLY#CAS-0001 y^2^ cho^2^ v^1^; attP40{nos-Cas9}/CyO. A line with a second chromosome landing site was obtained from Trent Perry, Bio21 Unimelb who had generated it using the Bloomington stocks #24749 (y[1] M{RFP[3xP3.PB] GFP[E.3xP3]=vas-int.Dm}ZH-2A w[*]; M{3xP3-RFP.attP}ZH-86Fb) and #25709 (y^1^ v^1^ P{y[+t7.7]=nos-phiC31\int.NLS}X; P{y[+t7.7]=CaryP} attP40) to generate second chromosome landing site flies (09w) in a *white* null background. We also generated our own w^1118^; attP40{nos-Cas9_NH} line by injecting 09w with a plasmid (pnos-Cas9-nos from Simon Bullock http://n2t.net/addgene:62208 ; RRID:Addgene_62208 (37)) to generate a nos-cas9 fly line that had the advantage of not requiring a second chromosome balancer.

### 2.3 Fly crosses and Homing

The exact crossing schemes are shown in the Supporting Information. The *UAS* mini-*Ace* transgenes, were combined with various *Gal4* driver lines, and were used to rescue *Ace* deletions (Figure S2). For the gene drives, which were designed to edit the native locus (Figure S3), we used a split drive design such that the nos-Cas9 would be provided on an unlinked site so that if, through misadventure, a fly were to escape from the laboratory the two components would segregate away from each other.

### 2.4 Malathion bioassays

Adult resistance to the organophosphate insecticide malathion was measured for the two transgenic mini-*Ace* overexpression lines PA-S UAS/nSyb-GAL4 and PA-R UAS/nSyb-GAL4, using 09w/nSyb-GAL4 line as control line. Adult flies were collected for two days, sorted by sex, and left for two days to mature. They were then transferred to 20mL glass scintillation vials that had been dosed with insecticide. For each scintillation vial 400μL of malathion solution (diluted in acetone) was added and distributed using a hotdog roller until the acetone had evaporated. Control vials were treated with 400μL of pure acetone. 15 single-sex flies were transferred into each vial without anaesthetisation, then the vials were sealed with cotton wool moistened with 10% sucrose solution. Experimental vials were left for 24 hrs at 25°C then mortality was scored. Malathion doses were tested by pilot assays and 5 doses plus control were used for the formal screen. Each dose was repeated three times.

### 2.5 Molecular analyses

Three-prime Rapid Amplification of cDNA ends was performed following the protocol of Scotto-Lavino *et al*.(38) using Phusion High Fidelity Polymerase (NEB) and the following two Gene Specific Primers: GSP1 (Ace-gD-F10): 5’-GGAGGGTTCATGACGGGTT-3’ and GSP2 (Ace-gD-F11):5’-GTCACCAGGGGCTTGGTAAAG-3’. For Quantitative Real time PCR, Trisure (Bioline) was used for RNA extraction, the GoScript Kit (Promega) was used to make cDNA and qRT-PCR performed on a BioRAD c1000 Touch thermocycler.

### 2.6 Conversion Efficiency

To assess the effectiveness of conversion we scored the frequency of dsRED among the progeny of single pair crosses of susceptibility gene drive (SGD) heterozygotes crossed to wildtype individuals of the opposite sex. As is convention, the efficiency (e) was calculated as double the deviation of the observed frequency of dsRED flies from the expected (0.5) frequency of dsRED flies, so that when there is no bias e=0 and when there is complete bias in favour of the marker e=1. Initially flies with the genotype w^1118^; nos-Cas9/+; SGD/+ were crossed to Raleigh line 804. Then the effect of Cas9 dosage was determined by scoring the progeny of flies that had the genotype w^1118^; nos-Cas9/nos-Cas9; SGD/+ crossed to w^1118^ homozygotes. To assess the effect of genetic background variation, SGD flies were crossed to particular DGRP flies so that they were heterozygous for SGD and a DGRP background and these were then backcrossed to flies from the same DGRP stock.

### 2.7 Population analyses

#### 2.7.1 Modelling expectation for cage experiments

Previously, one of us (B.J.C.) had developed a population based stochastic model of gene drives (39). Here we used a parameterization that reflected the cage experiments of our pre-zygotic gene drive: *Gene drive frequency* was initiated between 0.05 and 0.95, *Conversion efficiency* =0.25, *Population size* of 10^3^, we set *Selection coefficient* to 1 to represent the homozygous lethality of the *Ace* drive and the *Degree of dominance* to 0 and *Exposure rate* to 1 to code that lethality as recessive regardless of the environment. Each chromosome in the initial population could be targeted by the drive (there were no gene drive resistance alleles so *resistance level*=0) but a limitation of the model was that it did not allow gene drive resistance to increase over generations.

#### 2.7.2 Cage experiments

Three BugDorm-4S3030 cages (cubes with edges of 32.5cM) were used to conduct multigeneration gene drive experiments. 200 male and 200 virgin female flies all of which were heterozygotes for nos-Cas9 and the *Curly of Oster* (*CyO*) balancer chromosome were introduced into each cage. The *CyO* chromosome was necessary because the only second chromosome nos-Cas9 line we had available to us at the start of the experiment also had a lethal allele that prevented it from being homozygous. Eighty of the 200 males were heterozygous for the *Ace*-drive on the third chromosome, so that each of the three cages were seeded with the *Ace* drive marked by dsRED at a frequency of 10% (80 heterozygotes among 400 diploids). Six standard fly food bottles were placed in the cages with their lids off for 72 hours, so the flies had an opportunity to feed and lay eggs. The bottles were removed from the cages, cleared of flies, plugged, and kept at 25C. Two weeks later these bottles were placed back into the cage and unplugged to seed the next generation. The bottles were removed and replaced with fresh bottles and flies were left to lay for ∼three days (for the first three iterations) or ∼two days (as we controlled total number of flies by adjusting egg laying time). When the bottles with eggs were removed from the cage, the cage, including all the flies of the preceding generation, was placed into a chest freezer overnight and subsequently the dsRED frequency among these flies was counted. To sequence characterize individual non-dsRED flies from generation 10, PCR primers spanning intron 4 and flanking exons (CB_Ace_F2: ATCCCTGACATTTCCCATCA and CB_Ace_R3: GGAAATCCGCAGAACACGAC) were used where extension time was limited to 72C for 1 minute which would amplify an 866bp product across exon 4, intron 4 and exon 5 but not if the gene drive was in intron 4 (a full length dsRED and the sgRNA array would add about 2.7kb). Direct sequencing of PCR amplicons was performed using a Sanger sequencing service of Macrogen.

### 2.8 Testing the safety-switch

Simulations that model a population of sexually reproducing *Drosophila melanogaster* individuals occupying a two-dimensional rectangular (3:1) space with carrying capacity of 10, 000 were generated using SLiM version 3.6 (40) and were run on the University of Melbourne HPC Spartan using a shell script. The gene drive is released on the left-hand side of the space with the target subpopulation between the coordinates 0< x <1 while the non-target ‘shielded’ subpopulation lies within 2 < x < 3 also known as the shielded area. The migration between these two subpopulations occurs via an intermediate subpopulation where insecticide spray is simulated.

For each run, the population was initiated with 10,000 individuals. These individuals were assumed to be resistant to the insecticide. Each individual in the population was allocated a random position in the 2D space, The population follows an age structure with the overlapping generations occupying the same space.

#### 2.8.1 Movement

The individuals have a pseudo-random movement within the 2D space. In the initial simulations, this movement distance was drawn from a normal distribution with more than 99% individuals moving no more than 0.15. However, in latter simulations, the T-distribution was used to simulate the fat tailed dispersal kernels that correctly emulate *Drosophila* dispersal patterns (41,42) with some individuals moving well beyond the 0.15 limit. The mixture of normal model was used to emulate the T distribution with η=5 as SLiM does not contain T distribution as an in-built function. During the winter months, the movement distances were halved to emulate reduced dispersal in *Drosophila* populations.

#### 2.8.2 Reproduction

To simulate non-random mating, females are restricted to finding a mate within a certain radius dictated by the movement of individuals. After finding the mate, the model checks if both parents have sexually matured. If so, mating is regarded as successful. The number of offspring generated is dictated by the local spatial competition felt by the females. In overwintering drosophila populations, the fecundity was lowered.

#### 2.8.3 Mortality

The fitness and sexual maturity of an individual was tied to its age. Age based mortality is an essential part of the module that ensures elimination of the old, less fit adults from the population. This mortality is encoded as a sigmoid function fitted to developmental times and survival rates in the lab (43). During winter months, population parameters were altered to follow those of an overwintering study of *D. melanogaster* (44).

The drive was introduced near the left side of 2D space (x < 0.1) by converting a certain number of wild-type individuals into drive homozygotes. The drive was introduced in the 20th generation to ensure that the dynamics of the drive is not affected by the initial adaptation of the population to the given constraints. The drive allele assumes a representative homing Cas9 modification drive that converts insecticide resistant individuals into a susceptible phenotype, with a highly efficient Cas9 efficiency of 95%. To simulate the Cas9 activity with the drive allele, each generated offspring in that generation is checked for the presence of the drive and absence of Gene Drive Resistance allele. If the individual is heterozygous for the drive with no GDRs, the Cas9 makes the cut, converting it into either the drive allele or the GDR. For each individual, the probability of resistance arising is dictated by the resistance rate parameter that was set to 0.42 (45).

Since the drive allele renders the individuals carrying it to be susceptible, the spread of the drive can be restricted by spraying the insecticide between where the drive is present and the area to be shielded. The core principle of the control strategy is that the exposure to the insecticide would be lethal to the drive containing individuals, thus stopping the flow of the gene drive cassette into the shielded area. To simulate the safety-switch strategy, the insecticide was sprayed within a small width (W) of space present in the middle of the population after every generation is created. The mortality due to insecticide was drawn from a regression analysis of a malathion toxicological assay. Simulations were run for 3 different dosages: low, medium, & high. Each dose roughly corresponds to LD_50_, LD_75_, LD_100_. For each individual present within the spray window, the probability of exposure (IEP) dictates whether the focal individual is getting exposed. In the later stages of each generation, the shielded area is scanned for presence of the drive and its invasion into the wild population. The desired outcome of the insecticide barrier is to halt the spread of the drive into later parts of the rectangular space. The model checks the presence of the drive individuals in the x > 2.0. Presence of drive individuals beyond x > 2.0 signifies the failure of insecticide barrier control system. The probability of control was derived by calculating average successful control in 100 random seed simulations for each case.

The data was analysed and graphed using tidy-verse packages in R-studio to create heatmaps. The code is available at https://github.com/Charles-Robin-Lab/ControllingGeneDrives.

## Results

### 3.1 Mini-*Ace* rescues *Ace* CRISPR-knockouts

Knockout mutations of the acetylcholinesterase locus were created via CRISPR/Cas9 using four sgRNA’s (Table S1). Multiple deletions of 329, 462 and 600 nucleotides were recovered and none of the mutations could be homozygosed nor did they complement a deficiency line that lacked one copy of the *Ace* locus (BL-7974). These results confirmed that the CRISPR was targeting the *Ace* locus and that a functioning copy of acetylcholinesterase is essential for viability. Furthermore, analysis of the break points of these deletions suggests that all four sgRNAs can cut at the intended sites (Figure S1).

We then sought to rescue these CRISPR deletions with two ‘mini-*Ace*’ transgenes that encoded the PA isoform of acetylcholinesterase; one that encoded the organophosphate-susceptible allele of *Ace* (mini*ACE*-PA-S) while the other (mini*ACE*-PA-R) had four amino acids associated with insecticide resistance (I199V, G303A, F368Y, G406A). Both transgenes, were engineered to be under *UAS* control (36), and rescued *Ace* deletion alleles when driven with an *elav-Gal4* promoter (Figure 1; Figure S2). Furthermore, the transgenes were sufficient to maintain healthy ‘rescue’ stocks of *Ace* deletions for many generations (e.g. *elav-Gal4*; P{*UAS*-mini*Ace*-PA-R}/CyO; *Ace*^τ1^/ *Ace*^τ1^).

**Figure 1.**
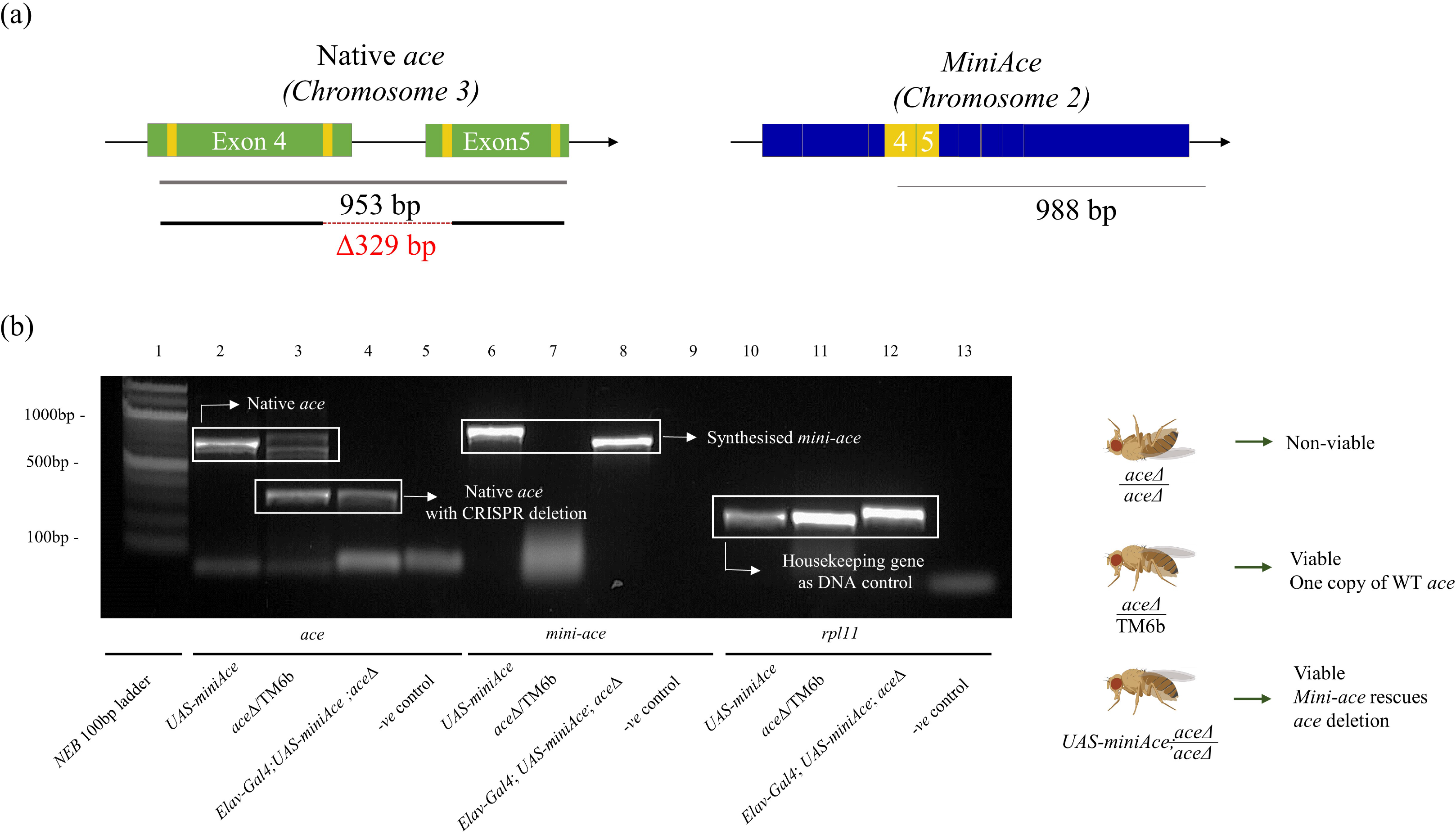
Mini-Ace construct (transgene of Ace coding sequence only) driven by *elav*-Gal4 rescues a CRISPR deletion within Ace. (a) The target sites of four sgRNAs in exons 4 and 5 of Ace (green) are represented in yellow and a CRISPR deletion between the two internal sgRNAs created a 329 deletion. The mini-Ace transgene encoding the full Ace-PA protein across ten exons yields a larger PCR fragment than the native Ace using different allele specific primers. (b) Agarose gel showing that a fly that is homozygous for the 329 deletion at the native Ace locus (lane4) survives because of the mini-ace transgene on chromosome 2 (lane 8).

### 3.2 Different forms of Mini-Ace transgene provide varying malathion resistance

Next, we examined the function of the mini-*Ace* transgenes with malathion assays. Ideally the transgenes would be regulated in a way that closely recapitulates native expression as that is less likely to introduce spurious fitness costs or gain-of-function phenotypes. However, we could not access an *Ace*-*Gal4* line and designing one proved challenging as the *Ace* promoter itself is poorly defined (46) and the small upstream intergenic region and the large first intron suggest that important regulatory elements may be contained within this intron. Analyses of single cell expression datasets of *Drosophila* brains show that the *NSyb* recapitulates the expression of *Ace* better than *elav* (Figure S4; 44). So, we drove expression of *UAS-mini-Ace* transgenes with the *NSyb-Gal4*. Adult flies, separated by sex, were tested for their ability to survive 24 hours of exposure to malathion. These experiments were done in the background of wildtype *Ace* alleles and unexpectedly the additional overexpression of mini-*Ace*-PA-S (as verified by Realtime PCR; Fig S5) did not significantly decrease mortality upon malathion exposure (LC_50_ of females of 0.44 vs 0.49 μg/vial: Figure 2). However, over-expression of the resistant form (PA-R) increased the LC_50_ by three-fold confirming that these amino acids substitutions do indeed contribute to malathion resistance in *Drosophila* (Figure 2; Table S2).

**Figure 2.**
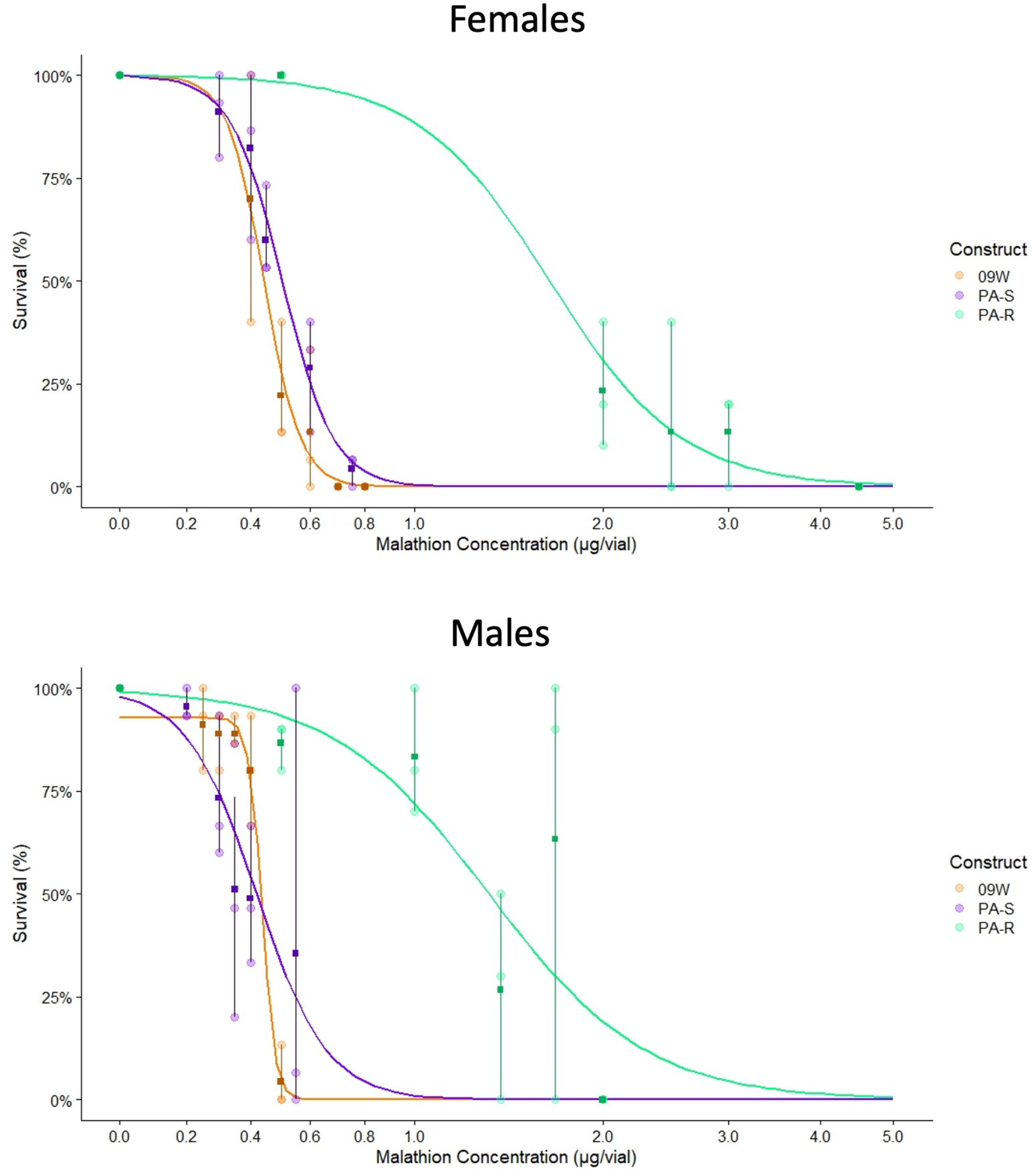
The mini-ace transgene carrying four known insecticide resistant variants is ∼3 times more resistant to malathion in both females (top) and males (below) than either of the mini-ace transgene with the four susceptible variants.

### 3.3 Design of a homing allelic drive that targets native exons

The three most common resistance mutations (I199V, G303A, F368Y) occur in adjacent exons (4 and 5) that are separated by an intron 159 nucleotides long. This provided the opportunity to design a homing gene drive in which the homology arms contained the ancestral susceptible forms and the intron encoded gene drive components – specifically the four sgRNAs and a dsRED marker gene (Figure 3). A plasmid construct composed of *Ace* homology arms and sgRNA elements (Supplementary Information S5) was initially injected into a line with a second chromosome *attP40* landing site. Once transgenic lines were established, constructs were homed to the natural *Ace* locus by crossing them to a line carrying nos-Cas9 on the X chromosome (Supplementary Information Figure S3). Five descendant lines had the homing to the 3^rd^ chromosome characterized using molecular markers. This revealed that three of these included all the desired features (Figure S6).

**Figure 3.**
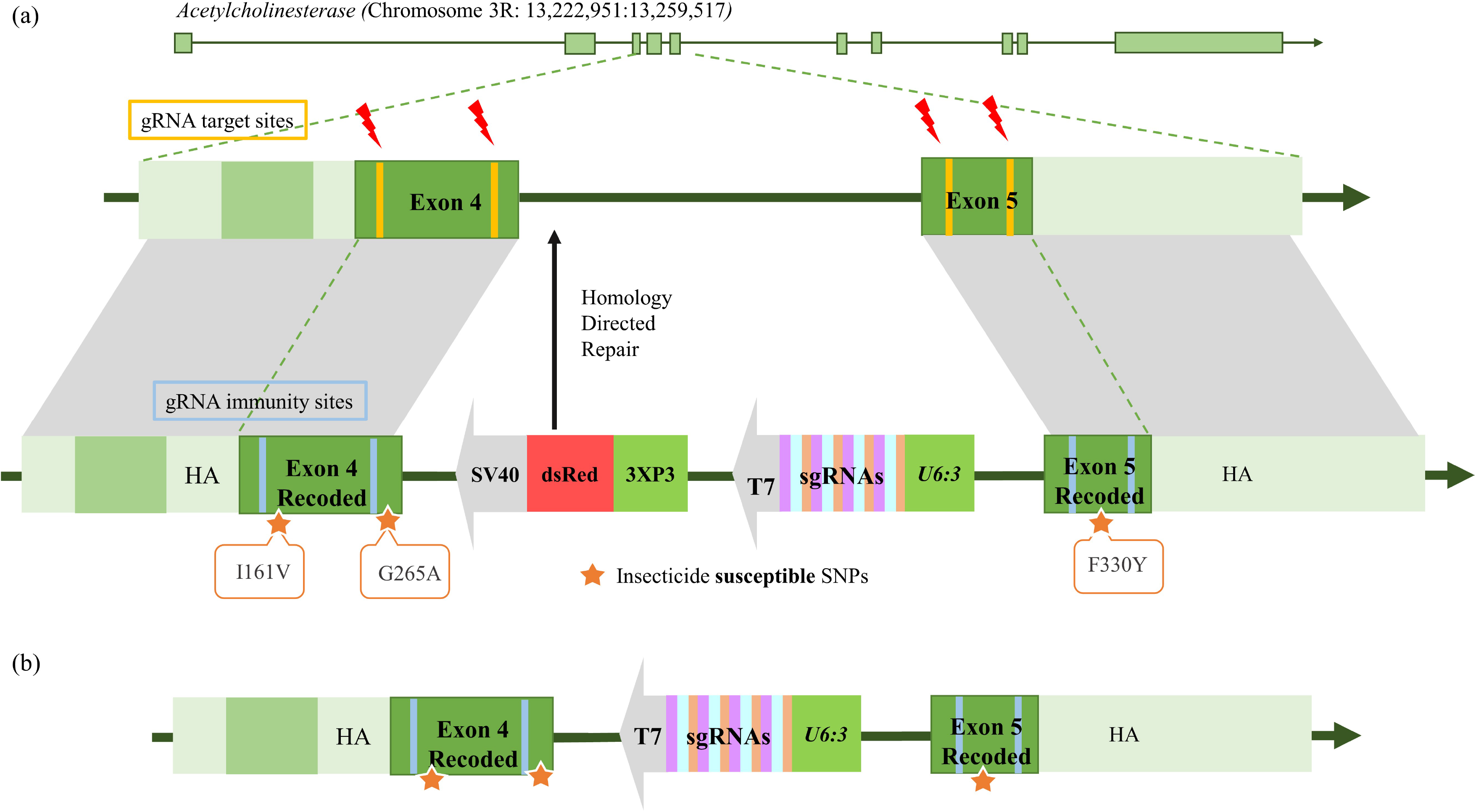
The susceptibility gene drive designed to replace three resistant mutations (I161V, G265A, F330Y) in homology arms that would not themselves be cut by sgRNAs because of ‘immunity’ changes (blue). The intron between Ace exons 4 and 5 had sgRNA/tRNA cassette encoded on the other strand. The construct represented in the middle panel also contains a dsRED marker gene (with 3xP3 promoter and SV40 terminator), whereas the construct at the bottom lacks the dsRED marker gene.

However, while we successfully ‘homed’ the construct to the natural *Ace* locus we were unable to homozygose the resultant drive alleles. Furthermore, the failure to complement *Ace* Deficiency lines, or *Ace* CRISPR generated alleles, indicated that the modifications that had been made at the *Ace* locus were not tolerated. The first time we could not homozygose the gene drive allele we discovered we had accidently introduced two non-synonymous mutations (N338K and H341Q) when making sgRNA immunity changes.

Correcting the errors and repeating the experiment produced the same result (the error had also been in the mini*Ace* constructs that rescued *Ace* deletions, so it was unlikely to be responsible for the homozygous lethal phenotype). Real-time PCR of heterozygous individuals using gene drive specific primers suggested that full length transcripts of the modified locus were not detectable. Rapid amplification of cDNA ends revealed that most transcripts ended prematurely within intron 4. In these transcripts, intron 4 had not been spliced out and a cryptic polyadenylation signal had putatively been used. In our third attempt, we removed dsRED and the cryptic polyadenylation signal, and we still could not get homozygous drive lines. It seems that placing the sgRNA construct into intron 4 of *Ace* was sufficient to destroy *Ace* function. However, the sgRNA’s themselves were expressed, they cut at the desired location, and enabled homing.

### 3.4 Genetic background and conversion efficiency

To quantify the extent of homing we initially crossed individuals heterozygous for the split gene drive features (nos-Cas9/CyO; SGD/+) to a wildtype stock (DGRP line Ral-804) and measured the proportion of flies bearing dsRED among the progeny. When the heterozygous parents were fathers, 947 of 1527 progeny had a dsRED phenotype (a conversion efficiency of 0.24; Tables S3). When they were females, 847/1380 progeny had dsRED eyes (conversion efficiency of 0.25). When we increased the dosage of Cas9 (nos-Cas9/nos Cas9; SGD/+) the efficiency increased to 0.38 (1349 dsRED flies out of 1959 progeny) although it is possible that this is associated with slight differences in genetic background rather than increased Cas9 dosage.

A previous study had shown that natural genetic variation encapsulated in the *Drosophila* Genetic Reference Panel (DGRP) altered conversion efficiency in a drive targeting the *yellow* locus (45). To see whether that genetic variation elicited a similar effect on our *Ace* drive we examined conversion efficiency among 13 DGRP lines mostly chosen from the extremes of the distribution found by Champer *et al.* (45). Two of these DGRP lines (Ral812 and Ral818) contained all three organophosphate resistance variants (199V, 303A, 368Y) while three only had the first one (199V, G303, F368; Table S3). In these experiments, flies heterozygous for a DGRP line and the split gene drive (nos-Cas9/DGRP-X; SGD/DGRP-X) were backcrossed to the DGRP line and the frequency of dsRED among progeny was scored. Ten replicate vials were set up for each of the 13 DGRP lines (Figure 4). This was done twice, with two different sources of 2^nd^ chromosome nos-Cas9. Two-hundred and forty of the 249 vials that produced offspring exhibited a bias towards dsRED. The total conversion efficiency of the two Cas9 sources was the same at 0.31 (3909/5955 flies and 4048/6168 flies). In each case the line effect was significant (One way ANOVA F=4.265 P<0.00001 and F=3.3277 P<0.00004). Generally, there was little correlation among the DGRP lines for the two Cas9 sources (R^2^=0.25, P=0.084) and no correlation with the order of DGRP lines observed by Champer *et al.* (2019) (R^2^=0.001, P=0.90). However, Ral181 had the lowest conversion efficiency in both Cas9 backgrounds and was among the ten lowest lines observed by Champer *et al.* (2019) and Ral812 had among the highest conversion efficiency in both Cas9 backgrounds and was the 11^th^ highest among the 138 lines assessed by Champer *et al.* 2019. Repeat assays of these two DGRP lines confirmed that the line effect was real. Only one DGRP line (Ral555), had a polymorphism at an sgRNA cut site (at 3R:13243715 twelve nucleotides from the PAM site in sgRNA 2; Supplementary Table S3) but it did not stand out from the other lines.

**Figure 4.**
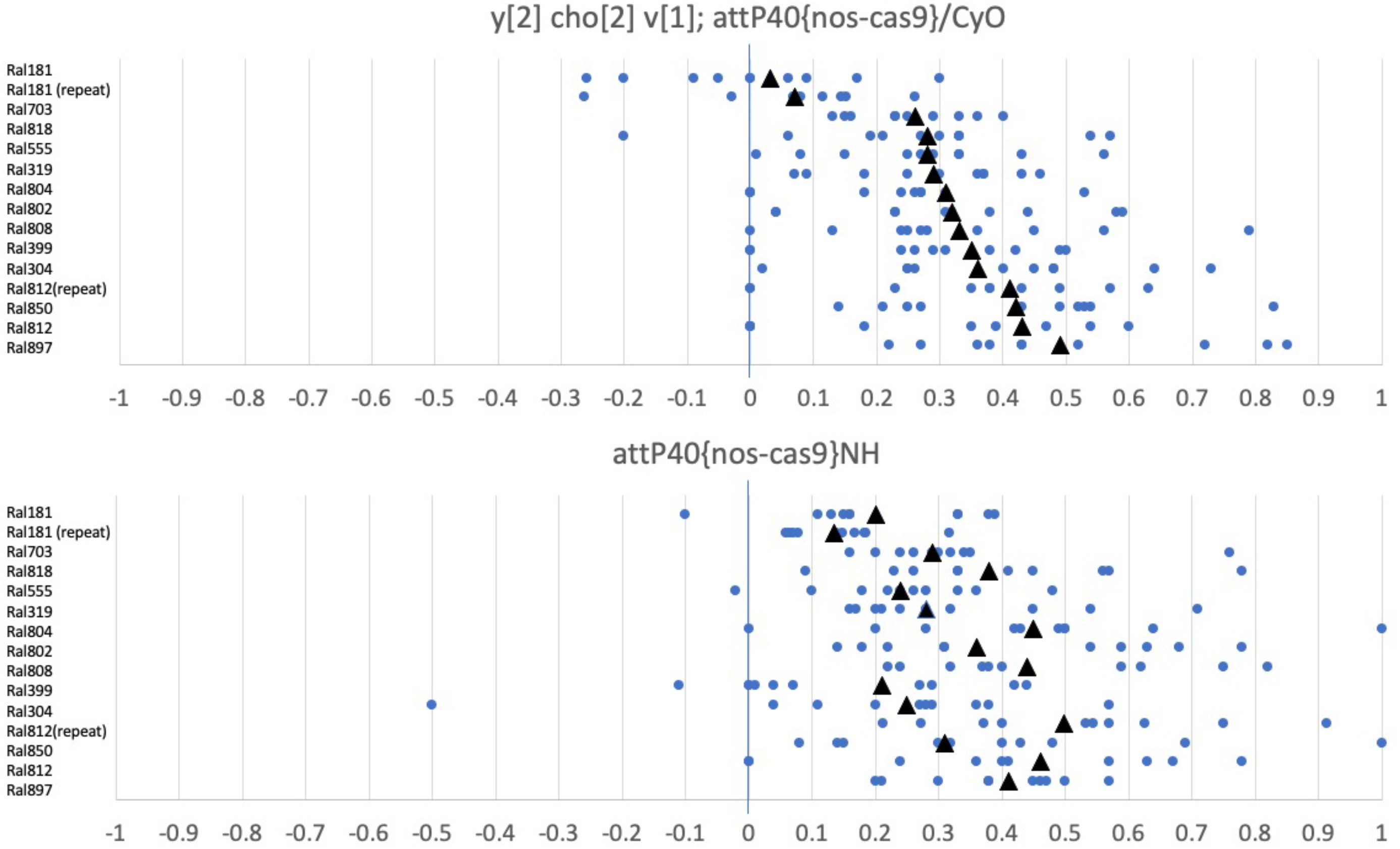
Inheritance bias and the variation in homing efficiency. The Ace gene drive was placed in various DGRP backgrounds and the bias in the inheritance of the marker (dsRED) was assessed (as twice the difference between the observed frequency and the expected Mendelian frequency). All genetics backgrounds showed bias towards the inheritance of dsRED indicating overall success of the homing strategy. Each blue dot represents a single vial containing the progeny of a single pair cross. The means of each background are shown as black triangles. Two different second chromosome nos-Cas9 stocks were used: (a) one from the y^2^ cho^2^ v^1^; attP40 {nos-Cas9}/CyO stock and (b) the other from a w^1118^; attP40 {nos-Cas9} stock. The lines of (a) are ordered by their average values and the lines of (b) are ordered as in (a). The measurements were repeated for the Ral181 and Ral812 to further establish the contribution of genetics to the efficiency variation.

### 3.5 Testing whether the *Ace*-drive acts as a toxin antidote drive in cage populations

As well as ‘homing’, the intron containing the sgRNA’s could act as ‘toxins’ that created lethal alleles in *Ace* when in a Cas9 background. As the drive chromosome had been engineered so that ‘immunity changes’ would prevent sgRNA’s cutting it, it can be thought of as bearing an ‘antidote’. Thus, the *Ace*-drive we had designed was also a ‘toxin: antidote’ drive which could increase its relative frequency by destroying wildtype alleles. We expected that among the non-dsRED progeny of *Ace* drive parents are those bearing alleles that have resulted from CRISPR/Cas9 double stranded DNA cleavage in at least some of the sgRNA targeted sites that were then repaired using the non-homologous end-joining (NHEJ) pathway. Such alleles are likely to disrupt the *Ace* gene and to be lethal and recessive. Thus, although the *Ace* SGD allele itself is homozygous lethal, it would also be capable of spreading non-drive lethal alleles. We wondered what the fate of populations bearing a drive such as this would be and so we performed simulations and established cage experiments.

Stochastic population-based gene drive simulations were used to model a pre-zygotic gene drive with a recessive lethal fitness cost. This showed that regardless of the starting frequency the gene drive converges on a frequency determined by the conversion efficiency (Figure 5a). We then tested this experimentally by establishing three population cages and examined the fate of dsRED marker over ten generations (Table S4). Each population cage started with 400 flies of equal sex ratio; each fly was heterozygous for a second chromosome bearing *nos*-Cas9 (with an unknown recessive lethal) and a CyO balancer chromosome. As both second chromosomes carried lethal alleles and complemented each other, and one chromosome was a balancer that prevented recombination, all flies in the multigenerational experiment would have curly wings and bear nos-Cas9. For each cage, 80 of the 200 initial male flies also bore the SGD on a third chromosome. In the first generation, rather surprisingly we observed the dsRED frequency to be 35% (1172/1805), 47% (761/1424) and 44% (943/1691) in the cages. If all the starting males had been equally reproductively successful and the conversion efficiency of the drive was 100% we would expect 40% of flies to carry the dsRED eye colour, and so it appears that, for unknown reasons, the dsRED males in the initial cross were more successful in mating (despite being the same age, reared on similar food, and genetic differences being limited to one third chromosome). From that high point, the frequency of flies with dsRED decreased over the ten generations to about 17% in each of the three cages (Table S4). Given all conversion efficiencies that we have measured above, bar one (Ral181), have been greater than 20% (Table S3), the allele frequency of the cages was lower than expected by the stochastic modelling. We hypothesized that this could be explained by gene drive resistance alleles arising. It is well established that when CRISPR/Cas9 double stranded DNA breaks are repaired by the NHEJ pathway, the motif targeted by the sgRNA is disrupted rendering the repaired chromosome immune to CRISPR/Cas9 in subsequent generations (48). We therefore sought to determine whether resistance was arising and if so whether it was through the NHEJ pathway.

**Figure 5.**
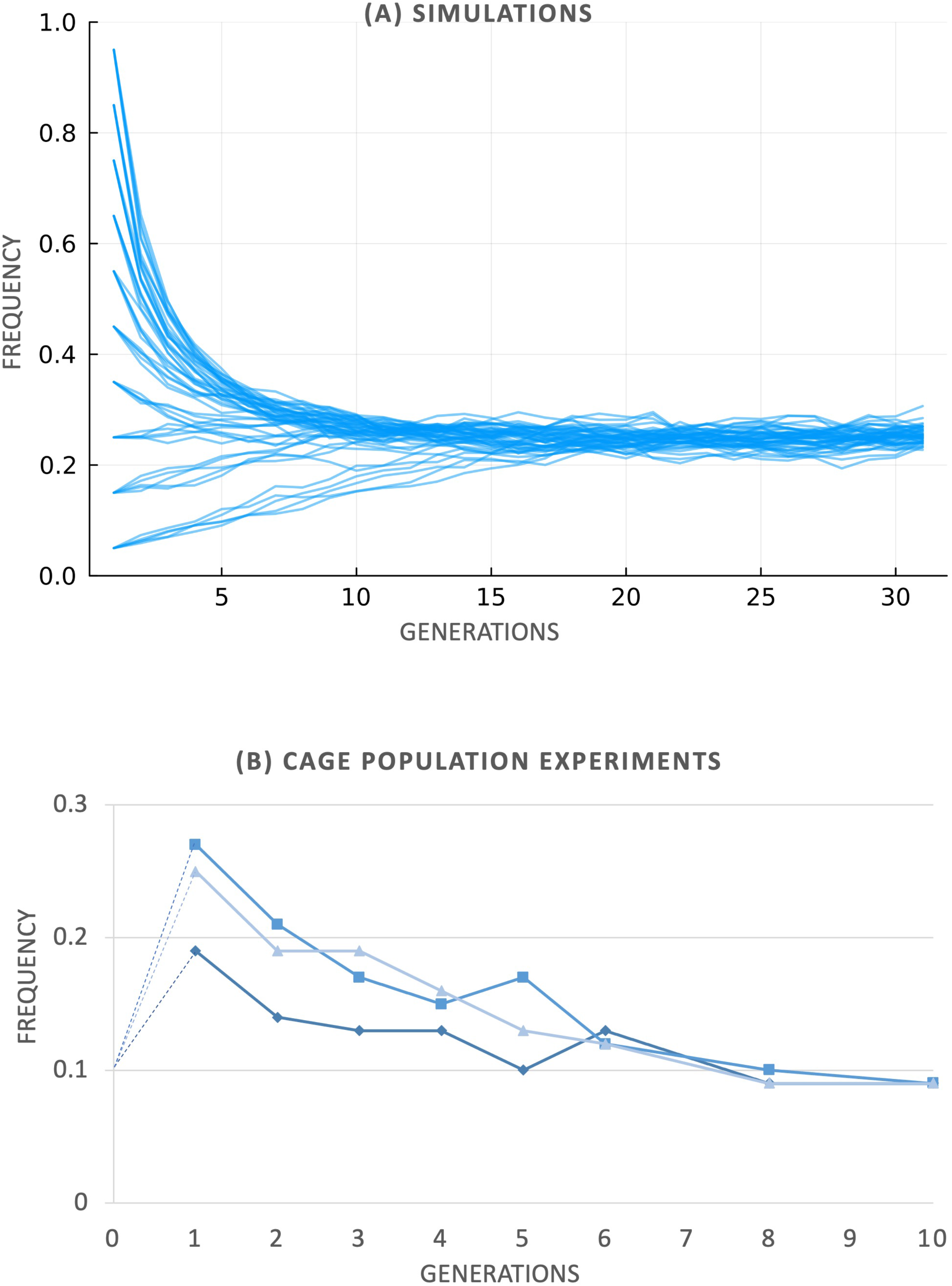
The fate of the ace-gene drive over generations. (a) Simulations of homozygous lethal gene drives with different starting frequencies (five replicates per starting frequency) in populations of 1000 individuals and a conversion frequency of 0.25 (b) The frequency of the gene drive (calculated as 1 – square root of the number of non-dsRED flies) in three caged populations each seeded with an initial frequency of 0.1 (Supp table 4).

### 3.6 Evidence of gene drive resistance

Virgins from the 11^th^ generation were extracted and mated in single pair crosses. Of 15 crosses involving a dsRED female and a wildtype male, only 6 were biased towards dsRED offspring. Of the 15 crosses involving a dsRED male and wildtype female only 8 were biased towards dsRED offspring and four produced no offspring at all. This is dramatically different to the 240/249 observed in prior experiments.

With such a low conversion efficiency, we sought to determine whether gene drive resistance had arisen over generations within the cages. We used PCR to amplify the *Ace* locus from non-dsRED individuals taken from the tenth generation of the caged populations. Agarose gels revealed that only one out of 60 individuals was heterozygotes for a detectable deletion. Unexpectedly, sequencing analysis from ten individuals revealed that seven had acquired at least some variants that we had introduced into the Susceptibility Gene Drive (SGD) construct (Figure 6). These individuals had sgRNA ‘immunity changes’ but lacked the dsRED and sgRNA cassettes suggesting that they arose via homologous repair and that the repair tracts were small.

**Figure 6.**
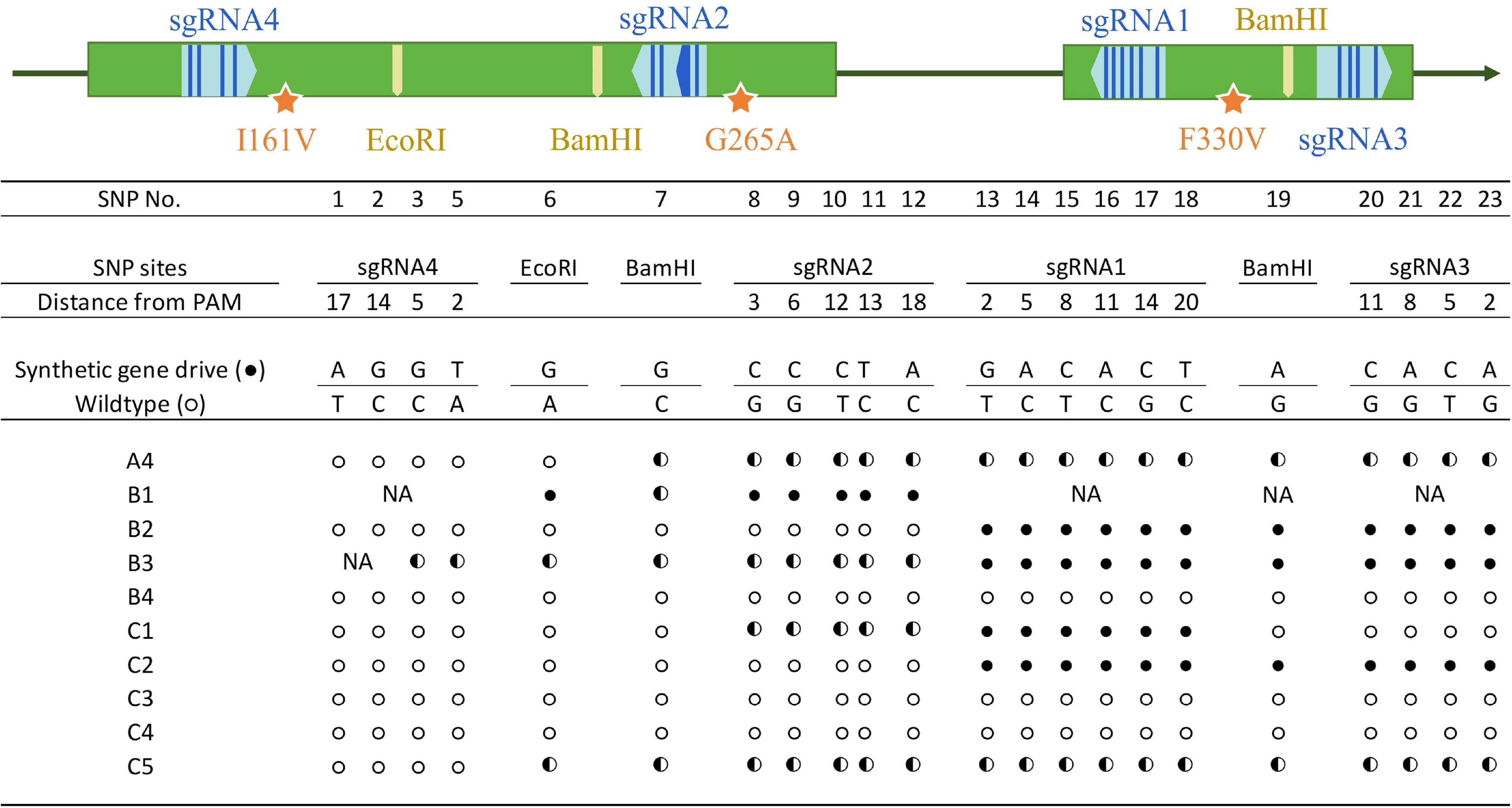
The acquisition of ‘immunity changes’ into non-drive chromosomes. The placement of four sgRNAs (pale blue arrows) on two adjacent exons of the Ace gene (green boxes). The amino acids conferring insecticide resistance are shown as stars and the sgRNA immunity changes represented by dark blue). Restriction enzyme sites that were deliberately modified in the synthetic DNA so they were different from wildtype Ace sequences are also shown. In total there are 23 sites where the synthetic construct had a state that differed to reference Ace sequence. The state at each of these 23 sites among ten lines isolated from the population cage at generation ten are shown as hollow circles (wildtype) or solid circles (synthetic by origin) or heterozygous (half:half circles).

In contrast, there were no novel mutations such as indels that would be expected if the NHEJ repair had been used. This suggests that mutation by NHEJ may only contribute a relatively minor amount to gene drive resistance observed in these experiments.

### 3.7 Could an insecticide be used to control unwanted spread of gene drives?

A central idea of insecticide susceptibility gene drives has been to revert insecticide resistance alleles back into susceptible ones so that insecticides can continue to be used (perhaps at a lower dose). A distinct and contrasting idea is that insecticide susceptibility could be engineered into gene drives (that are being used for other purposes – such as the suppression of local populations) so that the gene drives themselves could be prevented from spreading over time or space. By engineering chemical sensitivity into gene drives we could have a ‘safety-switch’ that could make gene drive technologies more controllable and palatable to regulators and the public.

To assess the feasibility of this strategy we simulated populations that inhabit a geographical area (of 3 x 1 arbitrary units) that was divided into three zones. In the first, the susceptibility gene drive was released, in the second insecticides were used to kill individuals bearing the gene drive, and the third contained a population that was to be shielded from the gene drive. If gene drive individuals made it into this third zone the ‘safety-switch’ strategy was deemed to have failed. In our simulations the gene drive spreads not only through the bias in inheritance but also by the movement of individuals bearing the drive, and that was initially modelled as following a normal distribution and then as a mixture of normal distributions. The width of the central zone, the spray window, was also varied between 0 and 1 units (Figure 7). We simulated the insecticide being sprayed once per generation and parameterized the proportion of individuals exposed to the insecticide in this window in a generation as a probability between 0 and 1 (Figure 7). These simulations showed that window width is irrelevant if the probability of insecticide exposure within the spray window is low (less than ∼0.2) and the probability of exposure is irrelevant if the window width is low (less than ∼0.2; the yellow L-shape of Figure 7) but otherwise the probability of control was high (especially with the wide spray windows). Also, the way that individual movement is modelled (normal dispersal versus T dispersal) affects the success of the control strategy, with wider spray windows being required to ensure population control under the more realistic fat-tailed dispersal distribution and the effect of insecticide dose also being lessened with such a dispersal kernel. In addition, we modelled the effect of seasonal population fluctuation on this strategy and found that if the total populations decreased due to winter, then gene drive control occurred more often.

**Figure 7.**
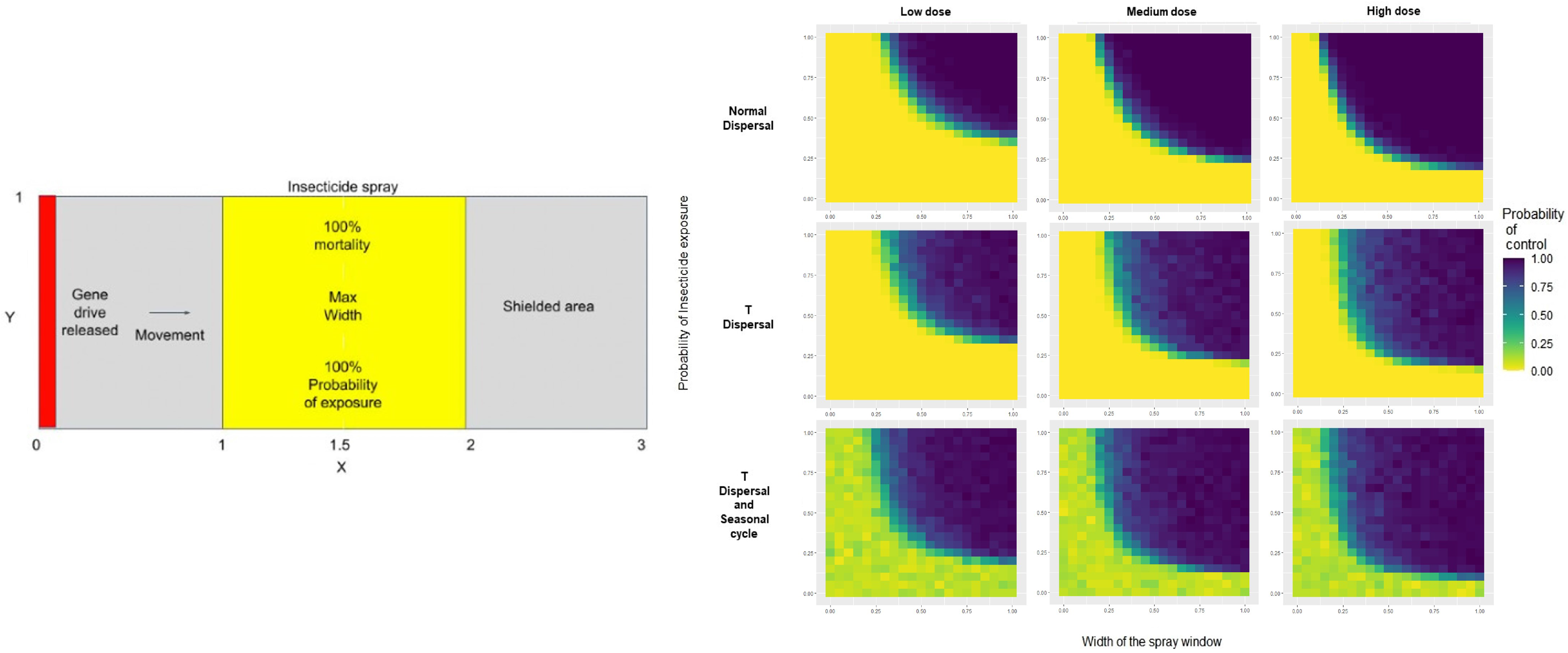
Modelling the safety switch in resistant *Drosophila* population (a) Schematic representation of the 2D space that harbours a resistant *Drosophila* population. This population can be divided into 3 subpopulations. First population (x<1) is the target population for the drive. The drive is released near x ∼ 0 (left side). Second subpopulation (x>2) is the non-target population. Insecticide is spray in the middle subpopulation (1<x<2). (b) Blue area represents the cases where gene drive was contained in the first population for different values of width of the spray window and probability of exposure. These simulations were repeated for different dosages (Low∼LD_50_, Medium∼ LD_75_, High∼ LD_100_) and different dispersal kernels and inclusion of seasonal cycle

## Discussion

Here we report an allelic susceptibility drive, in which machinery of the drive (four sgRNA’s and a dsRED marker that could be replaced by Cas9) was encoded in an intron of the acetylcholine esterase gene we were targeting. Such a design aimed to drive organophosphate and carbamate susceptibility through populations that had evolved resistance to them, reverting variants back to their ancestral susceptible state. We showed that homing did occur however it came at the expense of acetylcholine esterase function. The insertion of transgenic elements into intron 4 of *Ace*, seemed to prompt mis-splicing and result in the truncation of the *Ace* transcript. This discussion highlights some general learnings about gene drives, details some insights gained on acetylcholinesterase biology, and proposes some alternate susceptibility gene drive designs based on the *Ace* locus. We also comment on the feasibility and application of the ‘safety-switch’ concept that we have introduced.

### 4.1 Lessons on gene drive experiments

A key current limitation of CRISPR/Cas9 homing drives is the evolution of gene drive resistance (GDR). Our study alludes to distinct mechanisms of GDR. For homing drives GDR is thought to arise through the Non-Homologous End-Joining repair of double stranded breaks. Not only does the NHEJ pathway not copy the transgene by homologous recombination, but it also generates alleles that are resistant to further CRISPR/Cas9 cutting. GDR alleles have been classified into two types: r1 that do not alter the function of the targeted gene, and r2 that do, typically by generating frameshifting deletions or insertions (49). Building on this prior knowledge we implemented two ways to reduce gene drive resistance in our *Ace* susceptibility drive design (48). Firstly, multiple sgRNAs were encoded in an sgRNA/tRNA array (35), and so the odds of generating an r1 allele may seem remote because changes would have to occur at many independent sgRNA cut sites. Secondly, as *Ace* is a lethal gene, r2 alleles should be strongly selected against if exposed in a homozygous state or in combination with other *Ace* lethal alleles. Our cage experiment suggests that r2 alleles are arising with this drive, as we observed a large deletion that would obliterate sgRNA cut sites among the non-dsRED individuals in our cage experiment and we have confirmed without surprise, that most *Ace* mutations are lethal and so there will be strong selection against such r2 gene drive resistance alleles.

Of greater interest was the high frequency of r1 alleles that we uncovered in adult flies from our cage experiments (Figure 6). These suggest a novel mechanism by which GDR alleles can arise. Seventy percent of the flies we sequenced contain variants that we had engineered into the synthetic gene drive primarily so that the drive (which encoded sgRNA’s) did not cut itself. In these flies, variants had been transferred to a non-drive chromosome. It is improbable that these r1 alleles had arisen through an NHEJ pathway because their states agreed with what we engineered. We have two alternate explanations of how these alleles could come to be. The first is independent of sgRNA-mediated cleavage and that is the transfer of the immunity changes from the SGD allele to wildtype alleles occurred through recombination or gene conversion. This seems unlikely given the rates of recombination and gene conversion known in Drosophila (50). The second more intriguing possibility is that homologous repair of sgRNA-induced double-stranded break occurred using the SGD as a template. These patterns are consistent with a microhomology mediated repair mechanism perhaps via a synthesis-dependent strand annealing (SDSA) pathway (51) or the alternate end-joining pathway (52).

Regardless of mechanism, this type of resistance may be attributed to a flaw in our design. Other studies have heavily recoded their rescue alleles such that they are so divergent that they would not be used as templates in homologous repair (53,54). In contrast, we adopted a strategy of minimizing the changes in our target gene because in our judgement the heavy-recoding strategy risks subtly disrupting the function of the targeted locus in unanticipated ways that may introduce a fitness cost. However, in our strategy we inadvertently created templates that could be used by homologous repair pathways using short conversion tracts. Such a micro-homology mechanism could be of general importance because it could apply to naturally occurring polymorphisms that would not only increase in frequency because they are not cleaved but would additionally increase in frequency as they can be used as repair templates. Thus, GDR would arise rapidly with some gene drive designs.

Our data also indicates independence of cutting and repair among the four sgRNA’s we used. If multiple sgRNA’s had cut at the same time we would expect that large chunks of the *Ace* gene would be dropped out; and indeed, we did see such alleles (Figure S1). However, we recover flies in our cages that have altered only some of the gene drive targets suggesting that resistance to some is sufficient to increase the survival changes of the bearers. Presumably, this stems from the less than 100% cutting efficiency of any given sgRNA. This then suggests that naturally occurring polymorphisms that protects against single sgRNAs, could be selected for, and ultimately combine to give resistance to many sgRNAs in a multi-sgRNA gene drive.

Another type of GDR our data reveals is attributable to genetic background. Flies heterozygous for Ral181, an isogenic line from the Drosophila Genetic Refence Panel, repeatedly show little gene drive bias, whereas those heterozygous for Ral812 (that carries all the insecticide resistance mutations) show among the greatest bias in our experiments (conversion efficiency up to 50%). This cannot be attributed to polymorphism at the sgRNA cut sites as these lines have been sequenced and there are no polymorphisms in them.

Furthermore, these two lines also diverge in their conversion efficiency in the study of Champer *et al.* (45) who used a totally different CRISPR/Cas9 homing drive targeting the yellow locus. It is not clear whether the genetic differences are attributable to less sgRNA cutting (perhaps the background differentially modulates the *nos* promoter) or whether it can be attributed to relative preferences for different repair pathways (e.g. Homologous repair versus NHEJ). Champer *et al.* (45) suggests that there may be many polygenes of minor effect underlying the divergence between lines, but rare alleles in particular lines may also account for some differences, and the possibility of identifying a gene with large effect on conversion efficiency may motivate further characterization of these contrasting lines.

Finally, our the conversion efficiency of the *Ace* drive is low relative to those published (55–57). Perhaps this reflects the incorporation of SNPs into homology arms. However, our study does indicate that the efficiency could be improved with an increasing dosage or efficiency of Cas9. Thus increasing the dose of Cas9 would be recommended and likely could be further amplified following a recent finding demonstrated that Cas9 with two nuclear localization signals was much more efficient than Cas9 with only one (57).

### 4.2 Acetylcholinesterase biology

Like others before us we show that a transgene encoding *Ace* and lacking introns was sufficient to rescue *Ace* function (33). We note that such rescued flies cannot produce the rarer *Ace*-PC splice form, which has an alternate exon 10, encodes monomers (rather than dimers) that lack the GPI anchor. This suggests that the *Ace*-PC form does not have a vital function. We also confirm that the three amino acid substitutions I199V, G303A, and F368Y increase resistance to malathion, the last of which, is found in the original Morton and Singh (58) resistant line MH19 and has previously been shown to confer resistance through mini-*Ace* overexpression experiments (33). While Menozzi *et al.* (26) found that *in vitro* expressed acetyl cholinesterase with the I199V, G303A, F368Y enzyme to be 30 fold less inhibitable than the susceptible version of this enzyme, our *in vivo* experiments found the fold resistance at LD50 to be about three fold. Thus, other factors such as the half-life of the protein, the tissue of expression, and insecticide metabolism must be invoked to explain the 10x discrepancy.

Perhaps more inconsistent with earlier studies is the finding that extra expression of the susceptible form of Ace, as encoded by the mini-*Ace* construct, did not increase resistance. Previously Fournier *et al.* (1992) found that when the amount of Ace is reduced below wildtype (by examining heterozygotes and rescue constructs with low activity) resistance drops in a proportional way. They also report that natural occurring variation in *Ace* abundance is associated with insecticide resistance (31). Indeed, it is well established in *Anopheles gambiae* mosquito’s that copy number variation in the *Ace* gene yields organophosphate resistance, and low frequency gene copy number variation has been identified in *D. melanogaster* populations (59). However, in our toxicology experiments, mini-*Ace* was added on top of wild type *Ace*, raising the possibility that perhaps there is a maximum amount of Ace that can be expressed and have an impact on resistance. It also adds support to the proposition that that the duplications in other species may be adaptive, not because of any extra abundance of Ace, but because the duplication provides ‘permanent heterozygosity’ where the cost of an organophosphate resistance allele is offset by it being found with a wildtype alleles (30,60–62).

### 4.3 Better susceptibility designs

The intronic homing drive attempted here aimed to exploit the fact that insecticide resistance mutations were in adjacent exons. The targeted intron was small and while we targeted the part of it that was most variable across orthologs we appear to have disrupted the gene’s ability to function. This is despite genes commonly occurring within introns of other genes in *Drosophila melanogaster*, and that many natural and modified transposable elements occur within introns without egregiously disrupting gene function. However, our experience compels us to consider alternate susceptibility drives at the *Ace* locus.

One solution might be to put the drive machinery in another, larger intron, with the hope that larger introns might better tolerate insertions. We might also have greater success if we engineered intron splice sites so that they are stronger.

Another solution would be to build on the success of a mini-*Ace* rescue. SgRNA’s targeting *Ace* create lethal alleles and are therefore toxic, and a recoded mini-*Ace* could be considered as an antidote. There are two obvious initial drawbacks to such a design. Firstly, while the mini-*Ace* may rescue, it may still impose a cost relative to the wildtype alleles for which it would be competing in natural populations. Secondly, as *Ace* is a recessive lethal, selection will be slow to act as the toxic effect will occur in individuals homozygous for lethal alleles. Although perhaps a secondary drive mechanism, targeting a haplolethal genes as well could be used to spread the *Ace*-drive (54).

### 4.4 Chemical susceptibility as a safety-switch

A major concern of gene drive technology is their potential to spread into non-target populations. Several strategies have been proposed to deal with this (11,12,14,15) and here we explored the idea that a chemical sensitivity could be engineered into the gene drive so that insects bearing the gene drive could be differentially targeted by application of that chemical (10). There are many issues with this concept including the burden of engineering suitable chemical susceptibility genes as cargo into a gene drive. Our simulations also revealed that chemical exposure rate was an important parameter. Studies of insecticide application have revealed that many insects can exist within a ‘spray zone’ and yet they by chance avoid being directly exposed. Clearly such exposure rates vary dramatically with context (sprays from aircraft, booms, or by hand, or shipping container, or silo, and by crop) but we found high exposure rates would be necessary if there was any hope of this orthogonal method of controlling gene drives were to be achieved.

## Conclusions

As Ace is the molecular target of some of the more environmentally harmful insecticides a drive reverting resistant alleles to susceptible alleles may make it unsuitable for deployment in the field. However, we have explored the utility of *Ace* as a lab model for susceptibility gene drives. We found that it is possible to use a sub-gene homing design to replace key organophosphate resistance amino acid sites in acetylcholine esterase of *Drosophila melanogaster*. However, we were limited by the finding that the intron of *Ace* that we targeted could not tolerate transgene insertions. Nevertheless, the drive could bias inheritance regardless of the genetic background even though the genetic background may affect that bias. The efficiency of the drive also decreased over generations suggesting that gene drive resistance alleles accumulate easily especially through a previously unappreciated microhomology repair mechanism. Furthermore, through simulations we explored the idea that a chemical-based safety-switch could be used to limit the spread of gene drives and found that strategy would need to guarantee high chemical exposure rates and the movement of the targeted insects would need to be well characterized.

## Supporting information

Supplementary figure 3 Crosses for Homing

Supplementary figure 4 Expression like Ace

Supplementary figure 5 qRT-PCR

Supplementary figure 6 Evidence of homing

Supplementary Table 2 Dosage Mortality

Supplementary table 3 Conversion efficiency

Supplementary Table 4 Cage data

## Acknowledgements

We thank Max Scott and Jackson Champer for their discussions on gene drives, Clancy Lawler for his help with SLiM code and Ben Phillips for the suggestions about the T-dispersal function. This research was funded by the Australian Research Council (Grant number DP190102512). The research conducted here was licensed as a ‘Dealing Not Involving Intentional Release of GMOs into the Environment’ (Licence DNIR646 ‘Two types of split Drive for *D. melanogaster* lab experiments’) from the Office of The Gene Technology Regulator Australian Government.

