## Supplementary figures and images for "Acetylcholine esterase of *Drosophila melanogaster*: a laboratory model to explore applications of insecticide susceptibility gene drives"

### Supplementary Figure 1

#### Figure S1: Generation of Ace CRISPR deletions

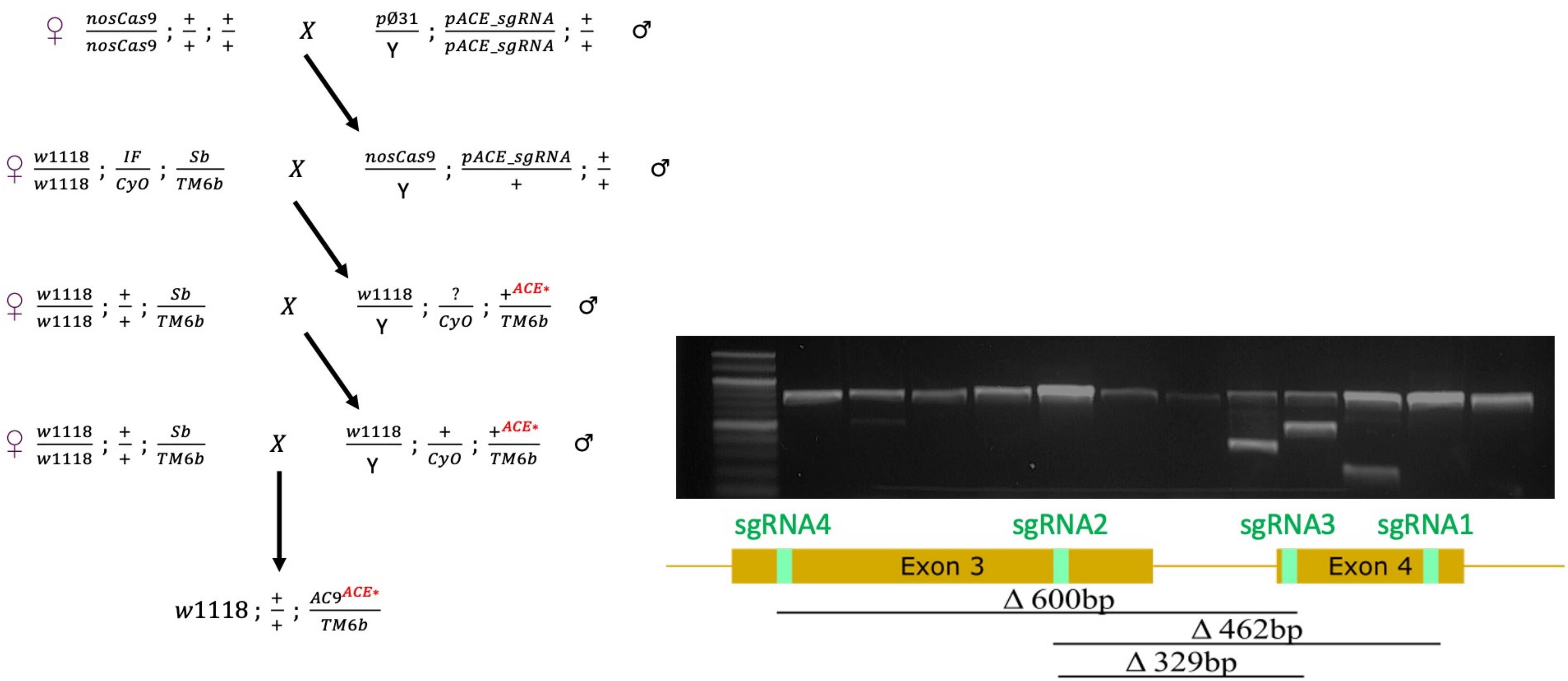

### Supplementary Figure 2

#### Figure S2: Mini-Ace rescues Ace deletions

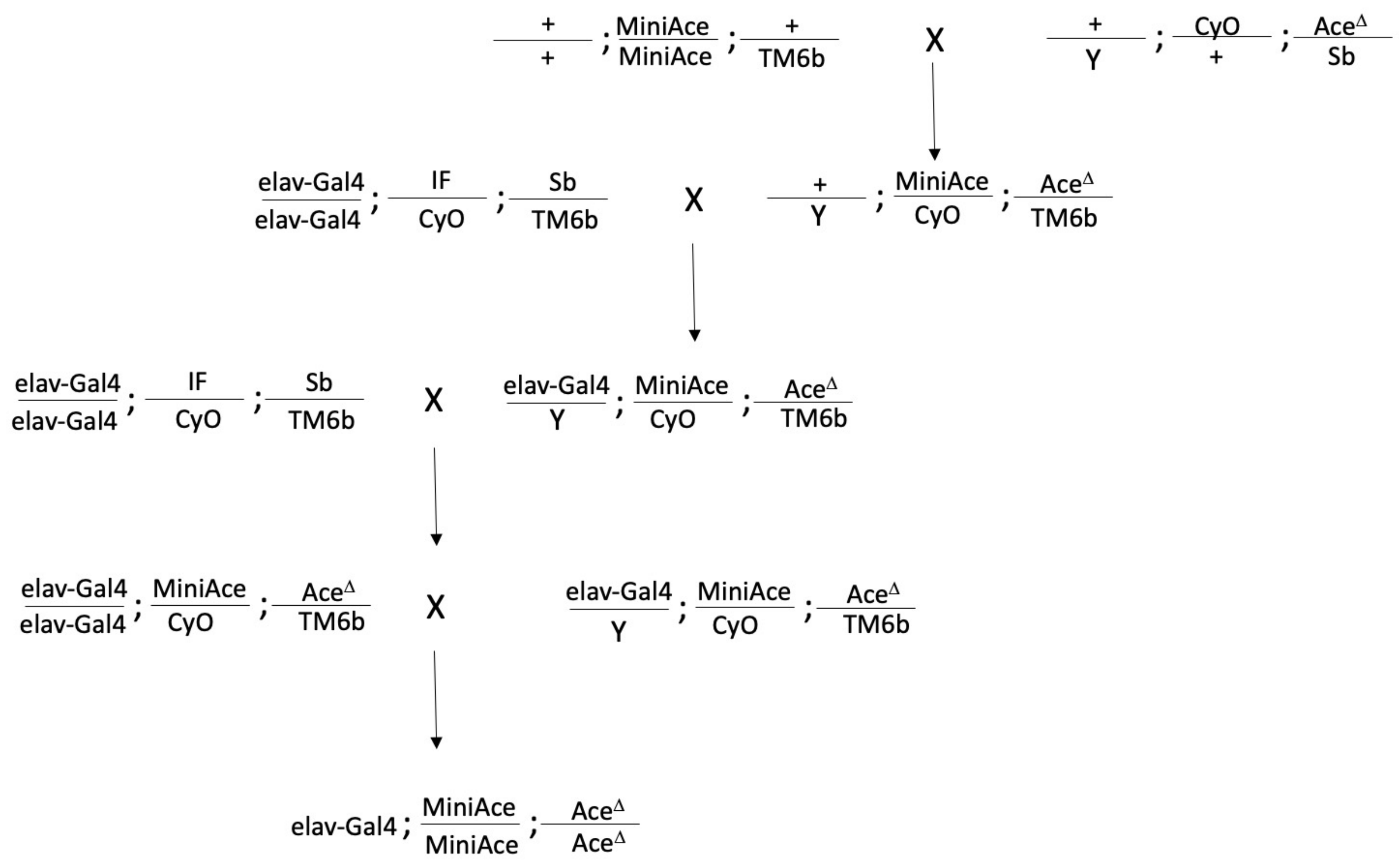

### Supplementary figure 3 Crosses for Homing

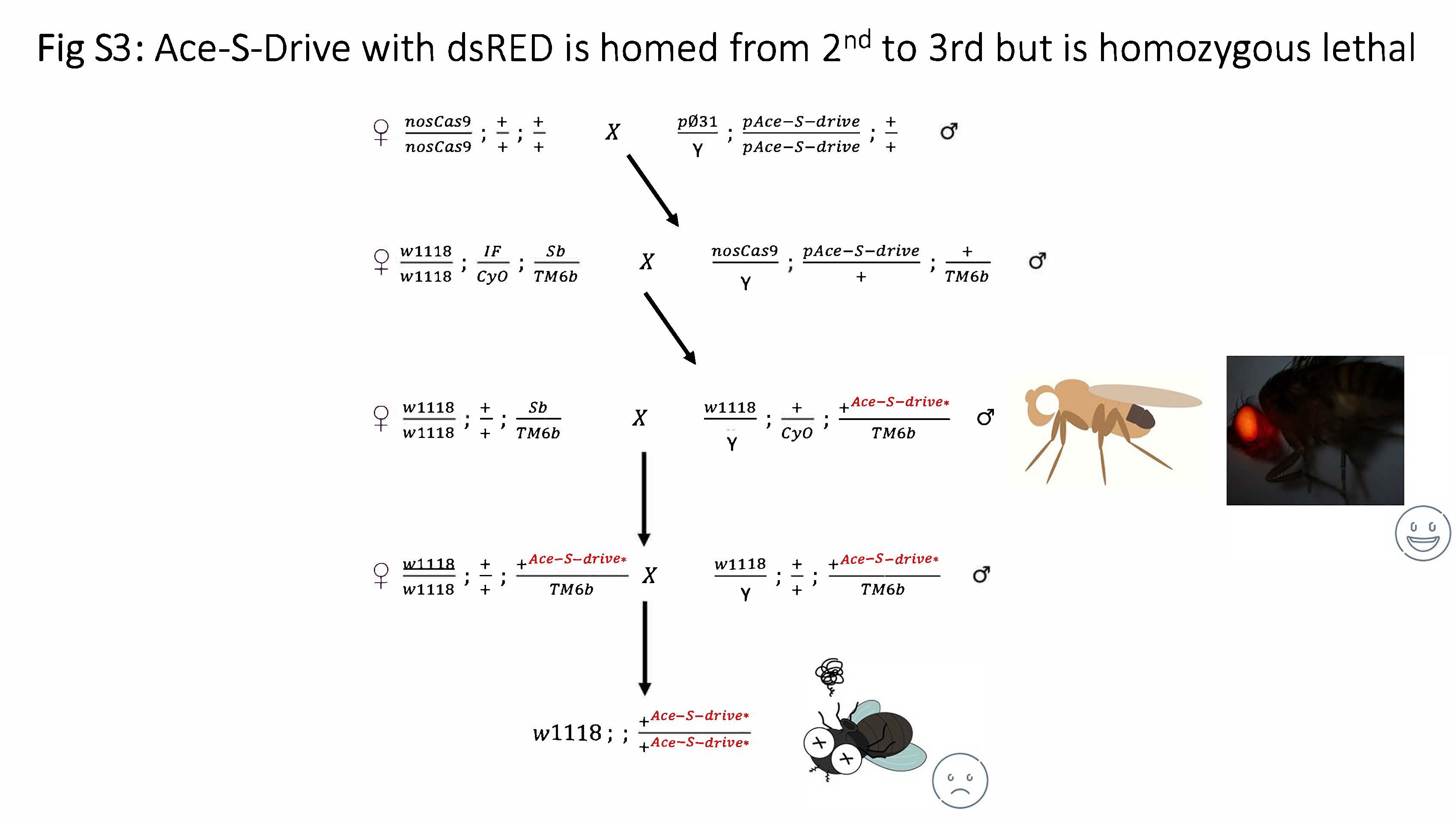

### Supplementary figure 4 Expression like Ace

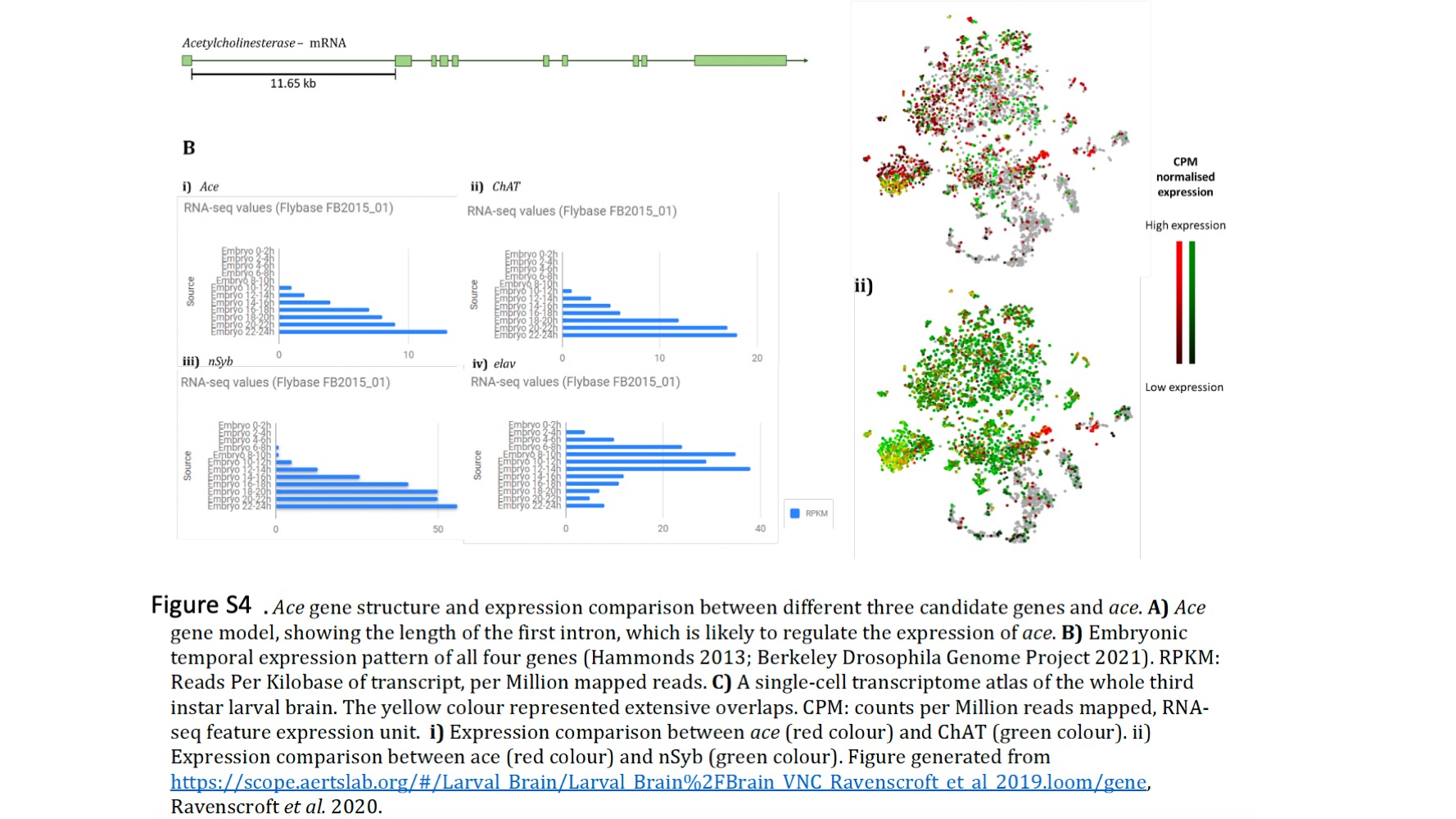

### Supplementary figure 5 qRT-PCR

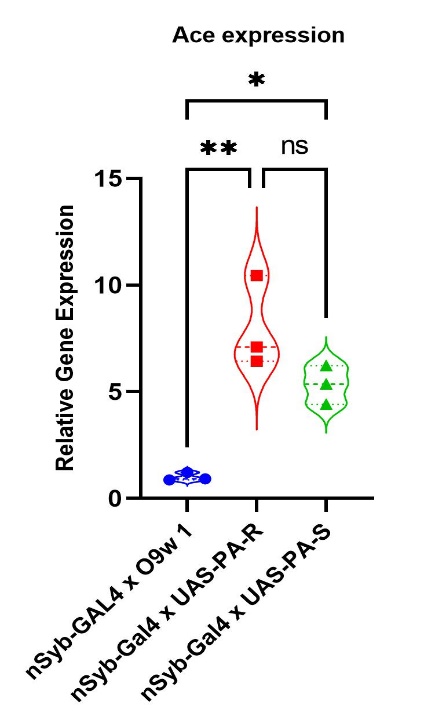
